# RNA isoform expression landscape of the human dorsal root ganglion (DRG) generated from long read sequencing

**DOI:** 10.1101/2023.10.28.564535

**Authors:** Asta Arendt-Tranholm, Juliet M. Mwirigi, Theodore J. Price

**Affiliations:** School of Behavioral and Brain Sciences, Department of Neuroscience and Center for Advanced Pain Studies, The University of Texas at Dallas, 800 W Campbell Rd, Richardson, Texas, 75080

**Author notes:** Corresponding Author: Theodore J. Price PhD Department of Neuroscience The University of Texas at Dallas 800 W Campbell Dr, BSB 14.102 Richardson TX, 75080.

**Keywords:** long read sequencing, dorsal root ganglion, splicing, hnRNPK, WNK1

## Abstract

Splicing is a post-transcriptional RNA processing mechanism that enhances genomic complexity by creating multiple isoforms from the same gene. Diversity in splicing in the mammalian nervous system is associated with neuronal development, synaptic function and plasticity, and is also associated with diseases of the nervous system ranging from neurodegeneration to chronic pain. We aimed to characterize the isoforms expressed in the human peripheral nervous system, with the goal of creating a resource to identify novel isoforms of functionally relevant genes associated with somatosensation and nociception. We used long read sequencing (LRS) to document isoform expression in the human dorsal root ganglia (hDRG) from 3 organ donors. Isoforms were validated *in silico* by confirming expression in hDRG short read sequencing (SRS) data from 3 independent organ donors. 19,547 isoforms of protein-coding genes were detected using LRS and validated with SRS and strict expression cutoffs. We identified 763 isoforms with at least one previously undescribed splice-junction. Previously unannotated isoforms of multiple pain-associated genes, including *ASIC3*, *MRGPRX1* and *HNRNPK* were identified. In the novel isoforms of *ASIC3*, a region comprising ∼35% of the 5’UTR was excised. In contrast, a novel splice-junction was utilized in isoforms of *MRGPRX1* to include an additional exon upstream of the start-codon, consequently adding a region to the 5’UTR. Novel isoforms of *HNRNPK* were identified which utilized previously unannotated splice-sites to both excise exon 14 and include a sequence in the 5’ end of exon 13. The insertion and deletion in the coding region was predicted to excise a serine-phosphorylation site favored by cdc2, and replace it with a tyrosine-phosphorylation site potentially phosphorylated by SRC. We also independently confirm a recently reported DRG-specific splicing event in WNK1 that gives insight into how painless peripheral neuropathy occurs when this gene is mutated. Our findings give a clear overview of mRNA isoform diversity in the hDRG obtained using LRS. Using this work as a foundation, an important next step will be to use LRS on hDRG tissues recovered from people with a history of chronic pain. This should enable identification of new drug targets and a better understanding of chronic pain that may involve aberrant splicing events.

## Introduction

RNA sequencing technologies have revolutionized our understanding of gene expression patterns in the human peripheral nervous system. Bulk RNA sequencing has provided a comprehensive insight into the transcriptome of the human dorsal root ganglia (DRG), including comparisons to other tissues and DRG in other species (Ray et al., 2018; Wangzhou et al., 2020). Furthermore, bulk sequencing of DRGs recovered from patients with pain have revealed how changes in gene expression may drive clinical phenotypes and how these underlying drivers differ in men and women (North et al., 2019; Ray et al., 2022). Spatial and single-cell technologies have given remarkable insight into the molecular makeup of human DRG (hDRG) neurons and other cell types in the DRG, revealing unique aspects of cell types and cellular makeup of the hDRG (Bhuiyan et al., 2023; Nguyen et al., 2021; Tavares-Ferreira et al., 2022; Yu et al., 2023). A major area that is still lacking is a careful examination of hDRG isoform expression. Some important progress has been made in this area as Sapio et al. have demonstrated how long read sequencing (LRS) can be used to characterize splicing in hDRG (Sapio et al., 2023).

Alternative splicing allows the generation of multiple isoforms from a single gene. This can affect protein structure, as well as expression patterns, through inclusion and exclusion of whole or partial exons, phosphorylation sites, or binding sites for RNA-binding proteins (Graveley, 2001; Stamm et al., 2005; Wang et al., 2015). Over 90% of genes in the human genome are estimated to undergo splicing, thereby enhancing diversity and complexity of the mammalian proteome (Pan et al., 2008; Wang et al., 2008). Tissue- and cell-specific alternative splicing programs drive inclusion and excision of sequences in transcripts expressed in neurons, driving development and associated with peripheral neuropathies (Lipscombe & Lopez Soto, 2019; Raj & Blencowe, 2015; Sapio et al., 2023; Su et al., 2021; Weyn-Vanhentenryck et al., 2018).

Functional implications of splicing associated with neuroplastic change and nociception have been shown in both rodent and human studies (Donaldson & Beazley-Long, 2016; Hulse et al., 2014; Liu et al., 2021; Song et al., 2023; Zhang et al., 2021). For instance, splicing of *VEGF-A* by SRPK1 has been shown to cause expression of two isoforms with contrasting pain-promoting (*VEGF-A_165_a*) or analgesic properties in mouse (*VEGF-A_165_b*) (Hulse et al., 2014). In a different example, Zhang et al showed alternative splicing of *HNRNPK* in the CNS of mice in opioid-induced hyperalgesia (OIH), by measuring expression of clusters of isoforms across conditions. *HNRNPK* exhibited significant alternative splicing in the nucleus accumbens associated with OIH, while a specific isoform (ENSMUST00000176849.7) exhibited lowered expressed in trigeminal ganglia (Zhang et al., 2021). *HNRNPK* encodes the RNA-binding protein hnRNP K, which has been associated with regulation of mu-opioid receptor (MOR) expression by promoting transcription and translation following morphine administration (Lee et al., 2014; Song et al., 2017). *OPRM1*, encoding the G-protein coupled receptor MOR, has additionally been shown to undergo splicing, facilitating extensive protein variation, with changes to number of transmembrane domains, consequently affecting opioid-induced signaling in humans (Donaldson & Beazley-Long, 2016; Liu et al., 2021; Xu et al., 2014).

Recently, results from a mouse model of inflammatory pain showed the downstream effect of splicing factor CWC22 on expression patterns of isoforms of secretary phosphatase or osteopontin, *Spp1* (Song et al., 2023). Following complete Freud’s adjuvant (CFA)-injection, an increase in *Spp1* variant 4 was observed and associated with a proinflammatory and pronociceptive mechanism. Despite evidence illustrating the importance of splicing as a functional mechanism of neuroplasticity and pain, our understanding of the isoform expression patterns in the hDRG is still lacking.

While isoform expression patterns can be inferred with SRS technologies, it has many limitations that can be overcome using LRS (Hu et al., 2021; Stark et al., 2019). Through optimized circular consensus sequencing, LRS provides a reliable insight into the transcriptome, potentially offering full-length transcript resolution (Sharon et al., 2013; Zeng et al., 2022). LRS has been used to characterize expression patterns of isoforms of annotated genes in human cells, an example of which is expression of transposon-derived short isoform of *IFNAR2* (Pasquesi et al., 2023). The short form IFNAR2 is associated with inhibited interferon signaling by forming a decoy receptor complex that does not produce intracellular signaling. The shorter isoform was found to be consistently more highly expressed than the long isoform in n human LRS datasets (Pasquesi et al., 2023). LRS also provides invaluable information to characterize RNA isoforms in the nervous system associated with neurodegenerative diseases as well as hereditary neuropathies (Sapio et al., 2023; Su et al., 2021). Sapio et al., used LRS and short read sequencing (SRS) to characterize *WNK1* isoforms, focusing on the HSN2 exon, wherein a mutation is associated with the development of hereditary sensory and autonomic neuropathy type 2 (HSNAII) (Shekarabi et al., 2008). An increase in the HSN2 expression was observed in the hDRG compared to the spinal cord (Sapio et al., 2023). Additionally, the majority of HSN2-expressing isoforms were found to contain a 27bp previously unannotated exon.

In this work, we utilized LRS to characterize isoform expression in hDRGs recovered from organ donors. Our aim was to create an overview of mRNA isoform diversity in the hDRG, providing a foundation for characterizing the role of alternative splicing in chronic pain development. We identified 19,547 coding isoforms, of which 763 isoforms utilized previously unannotated splice-sites. Furthermore, we independently identified a previously unannotated 27bp exon in *WNK1* in hDRG. Additionally, we verified the trend of comparatively increased short form *IFNAR2* expression in a human tissue not previously assessed. Finally, we present data evidencing novel isoforms of nociception-associated genes *ASIC3*, *MRGPRX1*, *SPP1* and *HNRNPK*.

## Materials and Methods

### Study design

The primary objective of this study was to characterize isoform expression patterns in hDRG. For this purpose, we performed LRS on lumbar hDRGs recovered from organ donors. DRGs from both male and female donors was included. All human tissue procurement procedures were approved by the Institutional Review Boards at the University of Texas at Dallas. For optimal sequence accuracy and variant calling Pacific Biosciences LRS technology was utilized (Su et al., 2021). *In silico* validation was performed by imposing strict cutoffs for confident detection of isoforms, in addition to verifying expression in a publicly available SRS dataset of hDRGs from independent organ donors (DBGap phs001158.v2.p1) (Ray et al., 2018). A formal pre hoc power analysis was not performed as no hDRG LRS data was available for estimation of isoform interindividual variabilities when we began this work. The analysis is considered a valuable overview of the isoform expression patterns, considering the inclusion of SRS data for validation. Gene names and isoform names are italicized to distinguish from protein names.

### Tissue procurement

Human DRGs used for RNA extraction were isolated from organ donors within 4 hours of cross-clamp (see donor demographics in **Figure 1A**). The tissue was frozen in dry ice immediately upon extraction and stored in -80°C freezer until the time of use.

**Figure 1:**
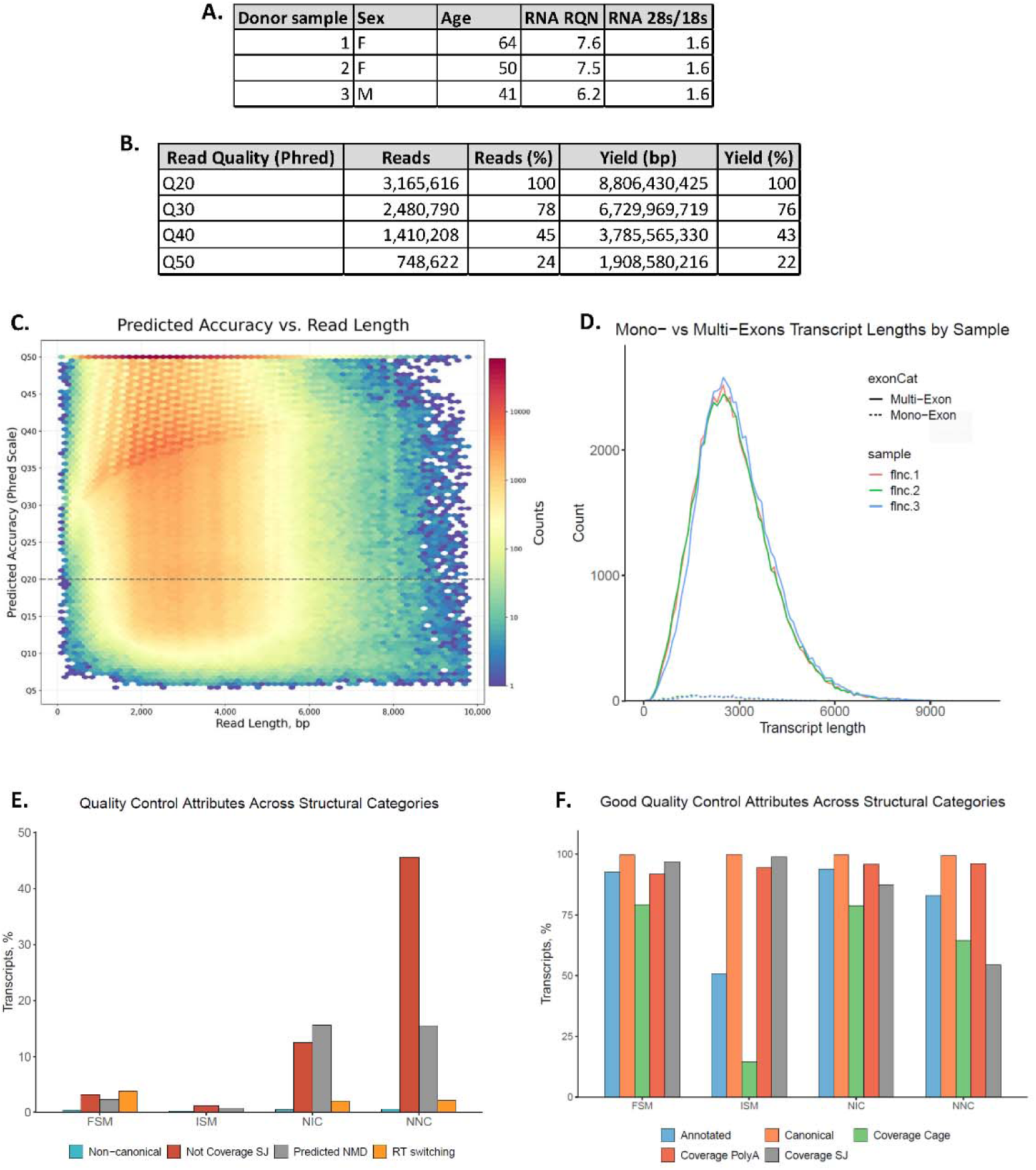
Descriptive characteristics and quality measures of LRS data. **A.** Donor demographics (sex and age) and RNA quality (RQN and 28s/18s) of hDRGs used for LRS. **B.** Table of read quality distribution of 3,165,616 reads show the number of reads and the bp yield grouped by estimated accuracy of bp measured as Phred score (Q20 = 99% to Q50 = 99.999% accuracy). **C.** Predicted accuracy of the bp call measured as Phred score compared to the length of reads. **D.** The number and length of mono- and multi-exonic reads for the 3 LRS samples showing comparable expression pattern with low interindividual variability. **E.** Quality control measures (usage of non-canonical splice-site, splice-junctions without coverage, predicted nonsense-mediated decay (NMD) and RT switching) are generally low for the 4 primary isoform subtypes (Full Splice Match, Incomplete Splice Match, Novel in catalog and Novel not in catalog), apart from usage of novel splice-junctions in novel isoforms as expected. **F.** Measures of good quality (annotated and canonical coverage, as well as coverage of poly-a motif, CAGE and splice-junction) were high for all 4 primary isoform subtypes, apart from coverage cage for ISM as expected. Figures were generated with SQANTI 3 (Pardo-Palacios, Arzalluz-Luque et al. 2023).

Human DRGs previously used for short read sequencing (SRS) were sourced from Anabios, see published methods in (Ray et al., 2018) for details.

### RNA extraction and sequencing

Total RNA isolation was performed from hDRGs of 3 organ donors using TRIzol according to the RNEasy Qiagen mini kit extraction protocol (see RNA quality in Figure 1A). Library preparation was carried out according to PacBio IsoSeq protocol with SMRTbell adapters to obtain a cDNA library of full-length transcripts. Circular consensus sequencing (CCS) was performed on the PacBio Sequel II equipment for 3plex on 1 SMRT Cell.

The SRS dataset was obtained from hDRGs from 3 female organ donors using Illumina TruSeq kits to generate polyA+ libraries used for 75bp paired-end sequencing (Ray et al., 2018).

### Isoform-level bioinformatic analysis

Only HiFi reads were included in analysis (CCS reads with Phred score greater than or equal to Q20). Using the IsoSeq3 pipeline, high quality full length non-concatemer (FLNC) reads were obtained by removing 5’ and 3’ primers, polyA tails and artificial adapters, followed by clustering and collapsing of redundant isoforms identified across samples using the GRCh38 reference genome.

Quality-control (QC) and filtering of the LRS-defined transcriptome of the hDRG was performed using the SQANTI3 tool-package (Pardo-Palacios et al., 2023; Tardaguila et al., 2018). QC was performed with standard settings according to the SQANTI3 protocol, as well as inclusion of CAGE-peak data and polyA list motifs for the GRCh38 genome. Publicly available SRS data from deeply sequenced hDRGs from independent organ donors, was added to the SQANTI3 Quality Check pipeline (DBGap phs001158.v2.p1) (Ray et al., 2018). An expression matrix for isoforms identified using LRS was generated by mapping the SRS-dataset to the LRS-defined transcriptome using kallisto. Splice-junction coverage was obtained using STAR (Ray et al., 2018). Filtering was performed using the SQANTI3 package with standard settings for rules-based filtering to target transcripts with unreliable 3’ regions related to intrapriming events, transcripts with RT-switching, and transcripts with all noncanonical splice-junctions and <3 counts for short read coverage. Isoforms that passed the initial filtering were further filtered by predicted coding potential, as well as cut-offs targeting transcripts identified in only 1 sample by LRS and with less than 2 transcripts per million (TPM) in SRS of hDRGs mapped to the LRS-defined transcriptome, facilitating an *in-silico* validation.

NetPhos3.1 was utilized to predict serine, threonine and tyrosine phosphorylation sites in neuronal networks, with kinase-specific predictions made for ATM, CKI, CKII, CaM-II, DNAPK, EGFR, GSK3, INSR, PKA, PKB, PKC, PKG, RSK, SRC, cdc2, cdk5, and p38 MAPK (Blom et al., 1999; Blom et al., 2004). IRESpy was used to predict internal ribosome entry sites (IRES) with an imposed prediction threshold of 0.1 (Wang & Gribskov, 2019). Structure and protein-folding of Swiss-Prot verified aa-sequences were explored using AlphaFold, and novel sequence folding was predicted using the Colab simplified version of AlphaFold v2.3.2 (Jumper et al., 2021).

## Results

### Long read sequencing facilitates isoform identification in hDRG

We used PacBio Sequel II IsoSeq technology to obtain an LRS-defined transcriptome of isoforms identified in hDRG. Following pre-processing, using the SMRT-link/IsoSeq pipeline, 3,165,616 reads with ∼8,8 billion bp were identified covering 152,264 isoforms (**Figure 1B**).

HiFi reads varied in length up to ∼10kbp with an average length of 2781bp, as expected of the human transcriptome (Lopes et al., 2021) (**Figure 1C**). High quality of sequencing was established, with a median Phred score of Q39 (99.99% accuracy), and 24% of reads ranked at Q50 (99.999% accuracy) (**Figure 1B, 1C**). The 3 hDRG samples sequenced were characterized by high concordance of transcript lengths and mono-versus multi-exon transcript expression, confirming similar library composition (**Figure 1D**).

SQANTI3 QC was performed to identify and characterize 8 structural categories of RNAs and their isoforms: Full Splice Match (FSM), Incomplete Splice Match (ISM), novel isoforms using canonical annotated splice-sites in a new combination (NIC), novel isoforms with at least 1 novel splice-site (NNC), Genic Genomic, Antisense, Fusion, and Intergenic. Over 90% of RNA isoforms were grouped into the first 4 categories. SQANTI3 furthermore facilitated quality check of the transcripts by assessing usage of non-canonical splice-sites, prediction of reverse transcriptase (RT) switching artifacts and nonsense mediated decay (NMD), as predictors of low quality. All 3 measures were shown to be low (**Figure 1E**). Usage of splice-junctions without coverage was observed primarily in novel isoform categories (NNC), as expected.

Furthermore, reference annotation, and canonical splice-site-, splice-junction- and polyA-site-usage, as well as SRS coverage were estimated to be high for all structural isoform categories, excluding inherent drop in ISM coverage (**Figure 1F**).

Through the inclusion of deeply sequenced SRS data of hDRGs from independent organ donors, filtering was performed using SQANTI3 to exclude isoforms predicted to be artifacts. Following this, isoforms of 14,056 unique genes were identified in the LRS samples. ∼70% (13,270 coding genes) of the protein-coding genes detected with SRS when mapped to the GRCh38 reference genome, were detected with LRS. 93,729 isoforms passed the initial filtering, including 39,421 FSM, 28,480 ISM and 25,413 novel isoforms (NIC or NNC), spanning 13,947 annotated genes and 109 entirely novel genes (**Figure 2A, 2B**). The isoforms identified in the hDRG were found to overwhelmingly use canonical splice-sites in both annotated and novel splice-junctions (**Figure 2C**). For less than 25% of genes only 1 isoform was identified, with most genes having more than 6 isoforms, showing expected high diversity of isoform expression patterns (Donaldson & Beazley-Long, 2016; Pan et al., 2008) (**Figure 2D**). Of the single-isoform genes, the majority were FSM with a transcript length slightly lower than the average (2535bp), including multiple *ZNF*-, *TMEM*- and *FAM*-genes. FSM were highly detected at all transcript lengths, while novel isoform discovery peaked at 4000bp length (**Figure 2E**). For further characterization, strict cutoffs were manually imposed to exclude isoforms identified in only 1 sample with LRS, as well as isoforms with less than 2 TPM from SRS mapped to the LRS-defined transcriptome (**Figure 2F**). 19,547 coding isoforms spanning 8256 genes were identified which passed the strict expression cutoffs (**Supplementary Table 1A**). 763 entirely novel isoforms containing at least one novel splice-junction were identified in the hDRG with LRS and the expression validated with SRS from independent human organ donors (**Supplementary Table 1B**).

**Figure 2:**
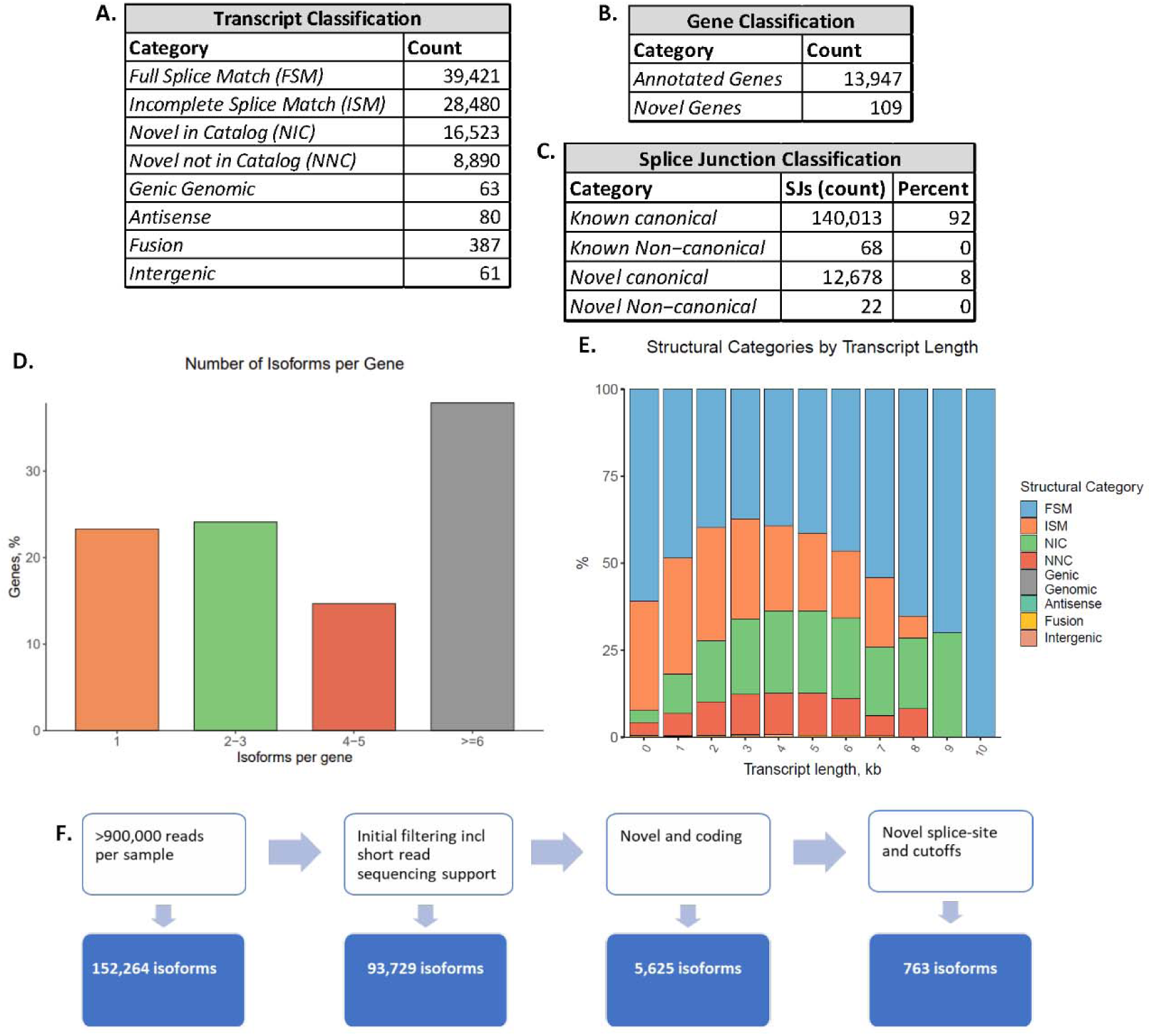
Analysis of LRS data. **A.** The number of transcripts identified with LRS and validated with SRS for each transcript category. **B.** The number of genes identified for each gene classification validated with SRS. **C.** The count and percent of splice-junctions categorized as known canonical, known non-canonical, novel canonical or novel non-canonical. **D.** The number of isoforms per gene expressed as percentages. **E.** The proportion of isoform classifications detected for each transcript length. **F.** Pathway of filtering from in silico validation using SRS to categorization by predicted coding potential, as well as novel splice-sites and expression cutoffs for both LRS and SRS ultimately identifies 763 novel coding isoforms. Figures 2D and 2E were generated with SQANTI 3 (Pardo-Palacios, Arzalluz-Luque et al. 2023).

### Distribution of gene expression with LRS reflects cell types in hDRG

Spatial transcriptomics, single-nucleus RNAseq, as well as extensive bulk sequencing of hDRGs, have previously provided a detailed image of the gene expression patterns expected to characterize the tissue at near single-cell type complexity (Nguyen et al., 2021; Ray et al., 2022; Tavares-Ferreira et al., 2022; Yu et al., 2023). To assess the gene expression of cell types expected in the hDRG, the isoforms were ranked according to LRS TPM. Of the 100 highest expressed isoforms, 28% were transcripts of genes corresponding to a panel of neuronally enriched markers of the hDRG (**Supplementary Table 1C**) (Ray et al., 2022). Furthermore, the highest expressed isoform was an FSM of the neuronal marker *NEFL*.

Further characterization of the data showed broad expression of genes enriched in neuronal subtypes identified with spatial transcriptomics and single nucleus sequencing from hDRG (Nguyen et al., 2021; Ray et al., 2018; Tavares-Ferreira et al., 2022). Isoforms of genes broadly enriched in neuronal cell-types were detected, including *SNAP25*, *TAC1*, *CALCA*, *CALCB*, *NEFH*, and *NEFL* (Figure 3A). Isoforms of genes expressed broadly in nociceptors were detected, including *NTRK1* and *SCN10A*. Furthermore, isoforms of genes enriched in several subtypes of C-fiber nociceptors, including cold-sensing nociceptors (*TRPM8* and *SCN10A*), silent nociceptors (*SCN11A*, *ASIC3* and *TRPV1*), pruritogen receptor-enriched nociceptors (*NPPB*, *GFRA2*, *IL31RA*, *TRPV1* and *SCN11A*), *PENK*+, and *TRPA1*+ nociceptors were detected. Additionally, isoforms of A-fiber proprioceptors (*PVALB*, *ASIC1* and *KCNS1*) were detected. While *SCN9A* has been confirmed to be expressed at the RNA level across sensory neuron subtypes, and functionally expressed in hDRG, the transcript was not detected at sufficient read-depth for our analysis (Li et al., 2018; Ray et al., 2018; Ray et al., 2023; Tavares-Ferreira et al., 2022). This is likely due to the transcript falling below the level of detection by LRS and the strict cutoffs we used. As observed in **Figure 3A**, of the 21 neuronal marker genes, 14 had more than 1 identified isoform (**Supplementary Table 2A**). For 13 genes, at least 1 FSM was identified, while novel isoforms were identified for 12 genes.

**Figure 3:**
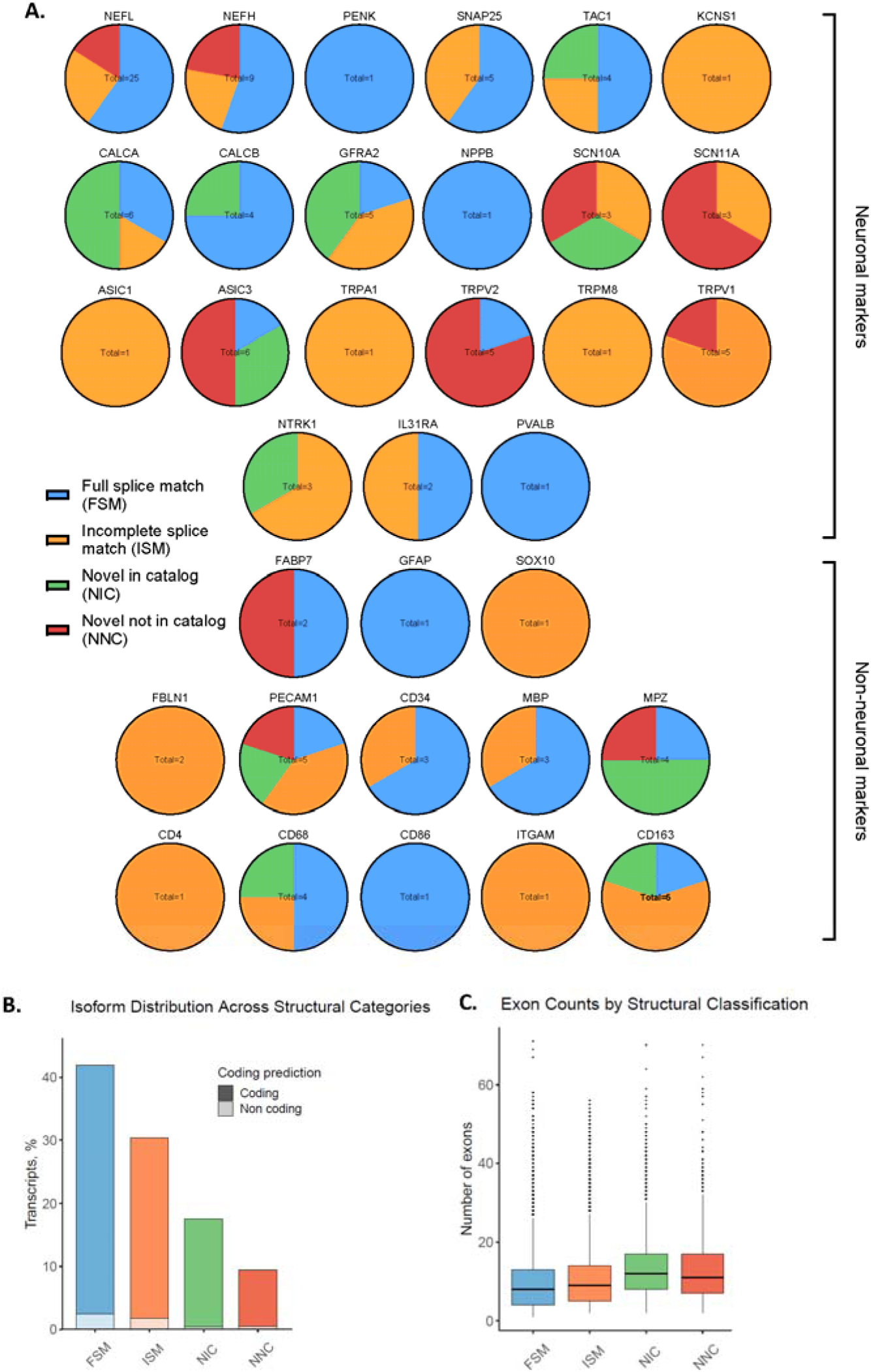
Structural variants of isoforms detected for neuronal and non-neuronal markers in hDRG. **A.** Isoforms of 21 neuronal markers and 13 non-neuronal markers detected with LRS and validated with SRS. **B.** The percent of coding and non-coding isoforms for the 4 primary structural categories. **C.** The average number of exons for the 4 primary structural categories. Figures 3B and 3C were generated with SQANTI 3 (Pardo-Palacios, Arzalluz-Luque et al. 2023).

Isoforms of genes enriched in non-neuronal cells were additionally observed, including broadly glial cells (*GFAP*, *SOX10*, *FABP7*), as well as specifically Schwann cells (*MBP*, *MPZ*), macrophages (*CD68*, *CD86*, *ITGAM*, *CD163*), and endothelial cells (*PECAM1*, *CD34*) (Xu et al., 2023). Lower expression was detected of isoforms of markers for fibroblasts (ISM isoforms for *FBLN1*, no isoforms identified for *COL1A2*), and T-cells (ISM isoforms for *CD4*, no isoforms detected for *CD8A*, *CD8B* and *CD3E*). Of the 13 non-neuronal markers, 7 had more than one isoform, an FSM isoform was identified for 9, and a novel isoform was identified in 5 (**Supplementary Table 2B**).

Broadly, a profile of gene expression patterns reflecting cell types expected in the hDRG was observed with LRS, with multiple isoforms detected for most genes. Of the 100 most highly expressed isoforms in the LRS-defined transcriptome, 98 were FSMs, 1 was an ISM (consortin, *CNST*) and 1 was a novel isoform (tumor protein D52, *TPD52*). Collectively, the LRS-defined transcriptome was characterized by a majority of coding isoforms across the structural categories, as well as consistent exon counts with a trend towards a minor increase in novel isoforms (**Figure 3B, 3C**). An increase in the coding region, provided by inclusion of additional exons, has the potential to have functional implications for the protein product (Raj & Blencowe, 2015; Stamm et al., 2005).

### Isoform expression patterns in the hDRG

The LRS-defined transcriptome was mined to assess detection and expression patterns of isoforms of genes of interest in hDRG. Published data has shown the shorter form of *IFNAR2* was more broadly expressed in human datasets (Pasquesi et al., 2023). This short form of the type 1 interferon receptor type 2 does not produce cellular signaling.

As illustrated in **Figure 4A** and **4B** only the short isoform of *IFNAR2* was identified in LRS-defined transcriptome of hDRG validated with SRS. The long isoform is detected following SQANTI3 QC and filtering, however, the expression level fails strict cutoffs. Consequently, the expression of *IFNAR2* isoforms in hDRG mimics the trend observed in previously tested human LRS datasets, with a higher expression of short form *IFNAR2*. An implication of this finding is that type 1 IFNs may signal primarily through IFNAR1 in hDRG, a hypothesis that can be tested in functional studies, as has been done previously in mouse DRG (Barragan-Iglesias et al., 2020).

**Figure 4:**
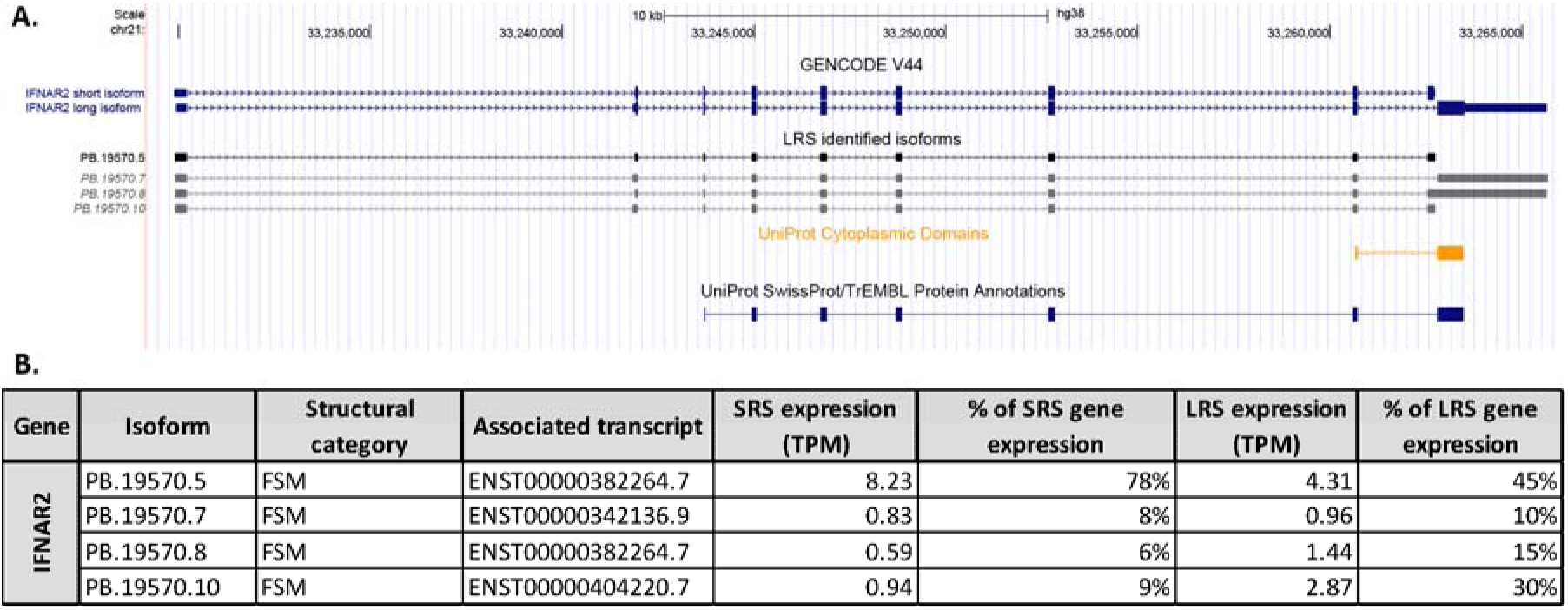
Isoforms of IFNAR2 detected with long read sequencing. A. The isoforms of IFNAR2 previously annotated in the GENCODE V44 library and the LRS-identified isoforms are shown. The italicized isoforms are identified but fall below the strict cutoffs imposed for detection. The UniProt protein and cytoplasmic domain annotation are shown. B. A table of the expression level (measured as TPM) for each isoform detected with LRS showing the percentage of total gene expression measured with SRS and LRS for each isoform.

LRS of hDRG have been utilized to characterize isoforms of *WNK1* (Sapio et al., 2023). A mutation in a DRG specific exon, HSN2, in *WNK1* is directly associated with the development of hereditary sensory and autonomic neuropathy type 2 (HSNAII) (Lischka et al., 2022).

Through characterizing *WNK1* isoforms in hDRG, a previously unannotated exon in *WNK1-204* (ENST00000530271.6), labelled 26c, was identified and found to be largely co-expressed with the HSN2 exon (Sapio et al., 2023). In our LRS-defined transcriptome of hDRG, 3 isoforms of the *WNK1* gene were identified.

One isoform utilized the previously unannotated splice-sites to include the 27bp 26c exon in the coding region (**Figure 5A**). The isoform containing the 26c exon, PB.12933.11, was identified in 2 out of 3 of the organ donors with LRS and made up 19% of the expression of the *WNK1* gene in the SRS dataset of different organ donors (**Figure 5B**). The additional 27bp sequence was predicted to add a 9 amino acid (aa) sequence to the coding region of MCPPAEPKS and creating a single aa deletion (aa2666, R). The structural layout of the folding of the protein produced from the isoform ENST00000530271.6 was found using AlphaFold, showing the insertion site of the novel exon at position aa2665 as shown in **Figure 5C**. Using NetPhos3.1, with ENST00000530271.6 as a framework, the additional coding exon was predicted to add a serine phosphorylation site, containing a potential consensus phosphorylation site for the kinase GSK3 (**Figure 5D, 5E**). Our data consequently confirms the presence of a previously unannotated 27bp exon in *WNK1* isoforms in the hDRG and suggests a potential unique function of this exon.

**Figure 5:**
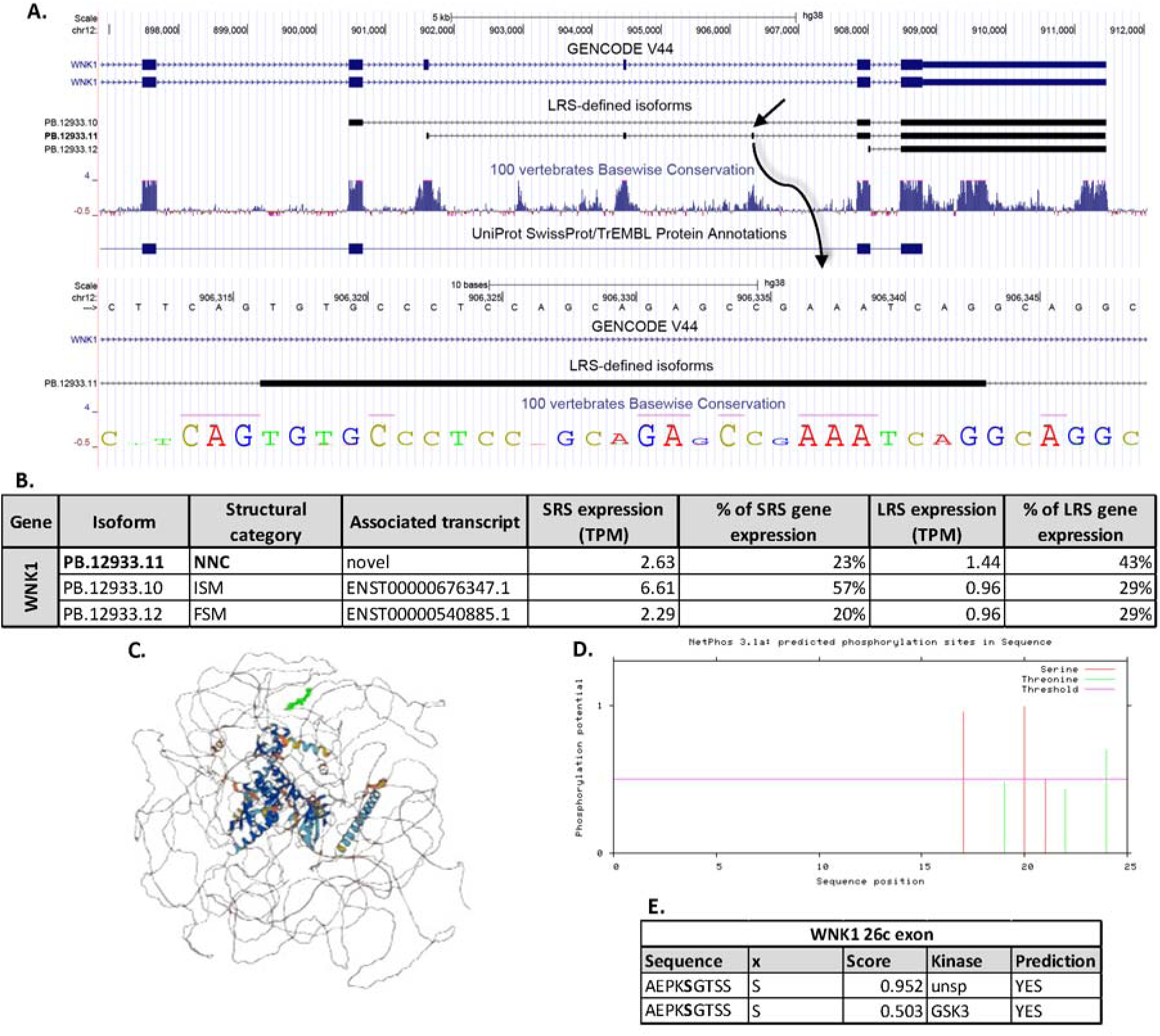
Isoforms of WNK1 detected with long read sequencing. **A.** The isoforms of WNK1 previously annotated in the GENCODE V44 library and the 3 LRS-identified isoforms of WNK1. The italicized isoforms are identified but fall below the strict cutoffs imposed for detection. The UniProt protein and cytoplasmic domain annotation are shown. A detailed look at the previously unannotated exon 26c is provided. **B.** A table of the expression level (measured as TPM) for each isoform detected with LRS showing the percentage of total gene expression measured with SRS and LRS for each isoform. **C.** Protein structure for the novel isoform, PB.12933.11, of WNK1 predicted with AlphaFold v2.3.2. The previously unannotated region, exon 26c, is highlighted in bold green. **D.** Predicted phosphorylation sites for serine and threonine are annotated in the novel exon 26c with NetPhos3.1. A cutoff of 0.5 for phosphorylation potential is imposed. **E.** A table of the predicted serine-phosphorylation sites in the WNK1 26c exon sequence shows a predicted GSK3 phosphorylation site.

In a CFA-induced mouse model of inflammatory pain, alternative splicing of *Spp1* caused an increase in *Spp1* variant 4 expression (Song et al., 2023). In hDRG *SPP1* isoform variant 1 (PB.5380.3) made up the primary expression detected using LRS (**Figure 6A, 6B**).

**Figure 6:**
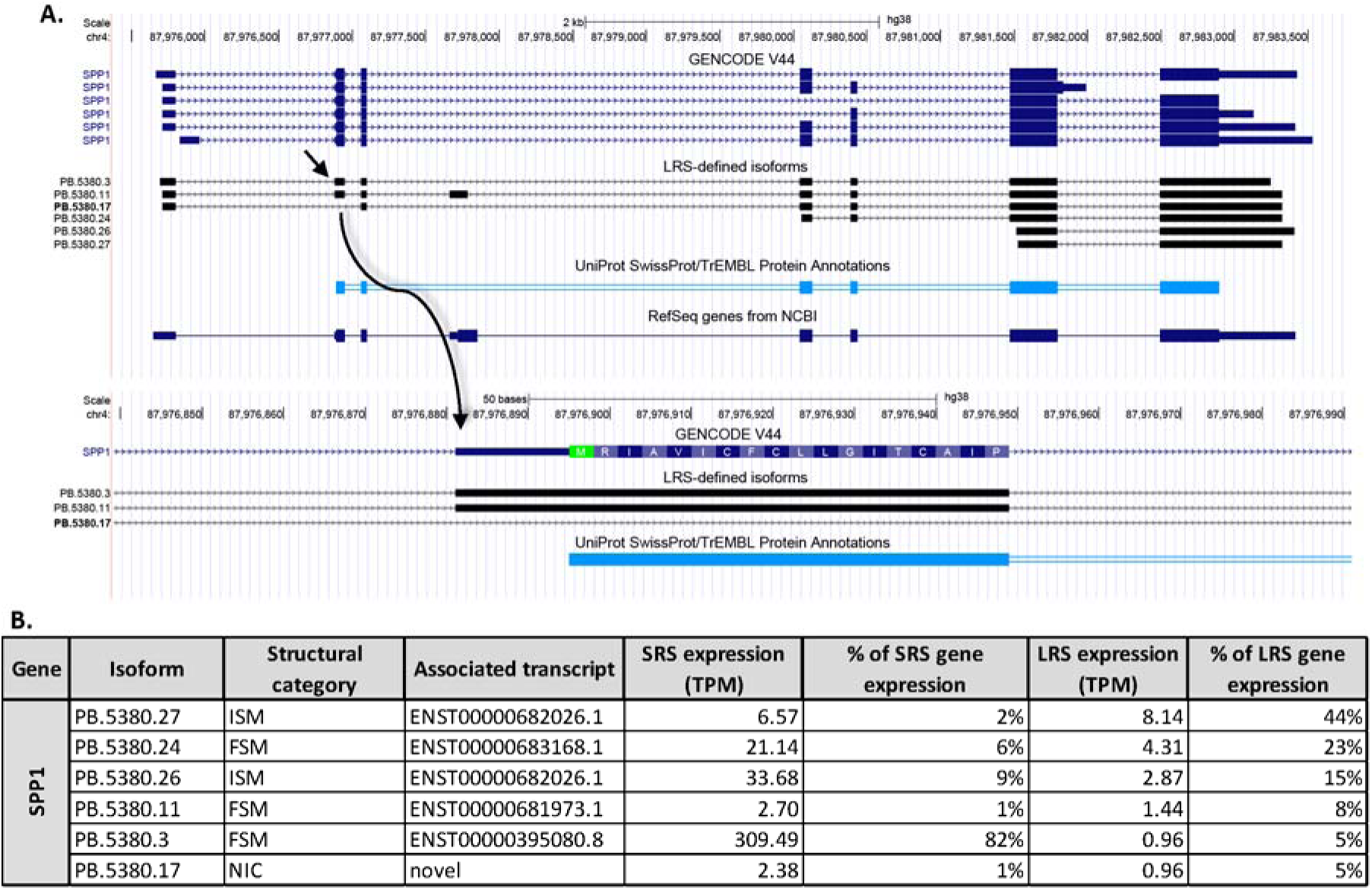
Isoforms of SPP1 detected with long read sequencing. **A.** The isoforms of SPP1 previously annotated in the GENCODE V44 library and the 6 LRS-identified isoforms. The isoform ID in bold indicates a novel isoform. The RefSeq sequence and the UniProt protein annotation are shown. A detailed look at the excised exon is shown, illustrating the localization of the start-codon, corresponding to the UniProt protein annotation. B. A table of the expression level (measured as TPM) for each isoform detected with LRS showing the percentage of total gene expression measured with SRS and LRS for each isoform.

These results could imply that expression of *SPP1* variant 4 may require alternative splicing induced by specific pain conditions in humans. Further studies of LRS on hDRGs from organ donors with a history of chronic pain is needed to address this. In the LRS-defined transcriptome, a novel isoform of *SPP1* was detected (PB.5387.17), which exhibited an excision of exon 2 (**Figure 6A, 6B**). Exon 2 contains the start codon, consequently the skipping of this exon may induce a disruption of the translation of the transcript. Further studies are needed to characterize the functional implication of expression of this isoform in hDRG.

### Novel isoforms are identified in the hDRG

When assessing only isoforms which pass the stringent cutoffs and with at least 1 novel splice-junction (NNC) as illustrated in **Figure 2F**, 763 entirely novel isoforms were identified in the hDRG (**Supplementary Table 1B**). Utilization of novel splice-sites facilitates the inclusion or exclusion of parts of the transcript. Of the 763 isoforms, multiple transcripts were from genes known to be involved in pain signaling including *ASIC3*, *MRGPRX1*, and *HNRNPK* (Donaldson & Beazley-Long, 2016; Liu et al., 2023; Zhang et al., 2021).

As shown in **Figure 7A**, 6 isoforms of *ASIC3* were identified, 3 of which were novel and utilized splice-junctions to exclude a region of 127 bp in the 5’-UTR. Sequences located in the 5’-UTR have previously been shown to drive post-transcriptional regulation and translation through inclusions of upstream open reading frames (uORFs), IRESs or binding sites for microRNAs (Ryczek et al., 2023). The excised region of 127bp excluded ∼35% of the total 5’UTR (127 of 355bp), including an upstream in-frame stop codon, annotated in the NCBI reference sequence at bp 197-199 (NM_004769.4). Using IRESpy (Wang & Gribskov, 2019; Yang et al., 2021), a potential IRES was predicted with a 0.1 prediction threshold for the canonical region which appeared to be retained despite the excision. The 3 isoforms were identified in hDRG from all 3 organ donors with LRS. Collectively, the 3 novel isoforms made up almost 50% of the expression of the *ASIC3* in the SRS dataset (**Figure 7B**). We were not able to identify any clear potential function of the excision from the 5’UTR of the ASIC3 transcript. Further work will be needed to understand the implication of this NNC.

**Figure 7:**
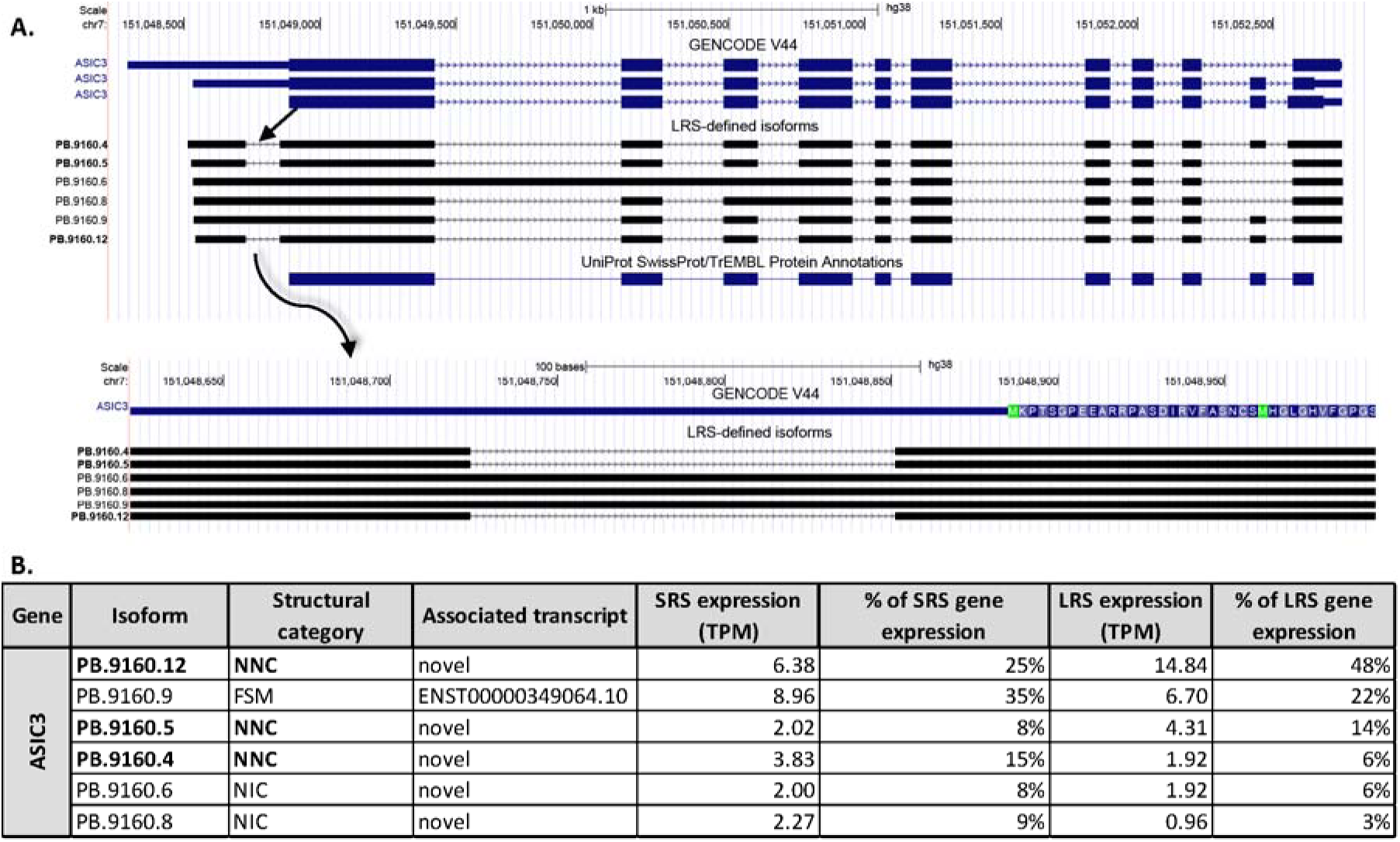
Isoforms of ASIC3 detected with long read sequencing. **A.** The isoforms of ASIC3 previously annotated in the GENCODE V44 library and the 6 LRS-identified ASIC3 isoforms. The isoform ID in bold indicates a novel isoform. The UniProt protein annotation is additionally shown. A detailed look at the excised region in the 5’ UTR illustrates the localization of the start-codon. **B.** A table of the expression level (measured as TPM) for each isoform detected with LRS showing the percentage of total gene expression measured with SRS and LRS for each isoform.

In contrast to *ASIC3*, where a novel splice-junction excised a region, an isoform of *MRGPRX1* was identified in the LRS dataset which utilizes previously unannotated splice-sites to include sequences in the transcripts. Four isoforms of *MRGPRX1* were identified in hDRG, shown in **Figure 8A**.

**Figure 8:**
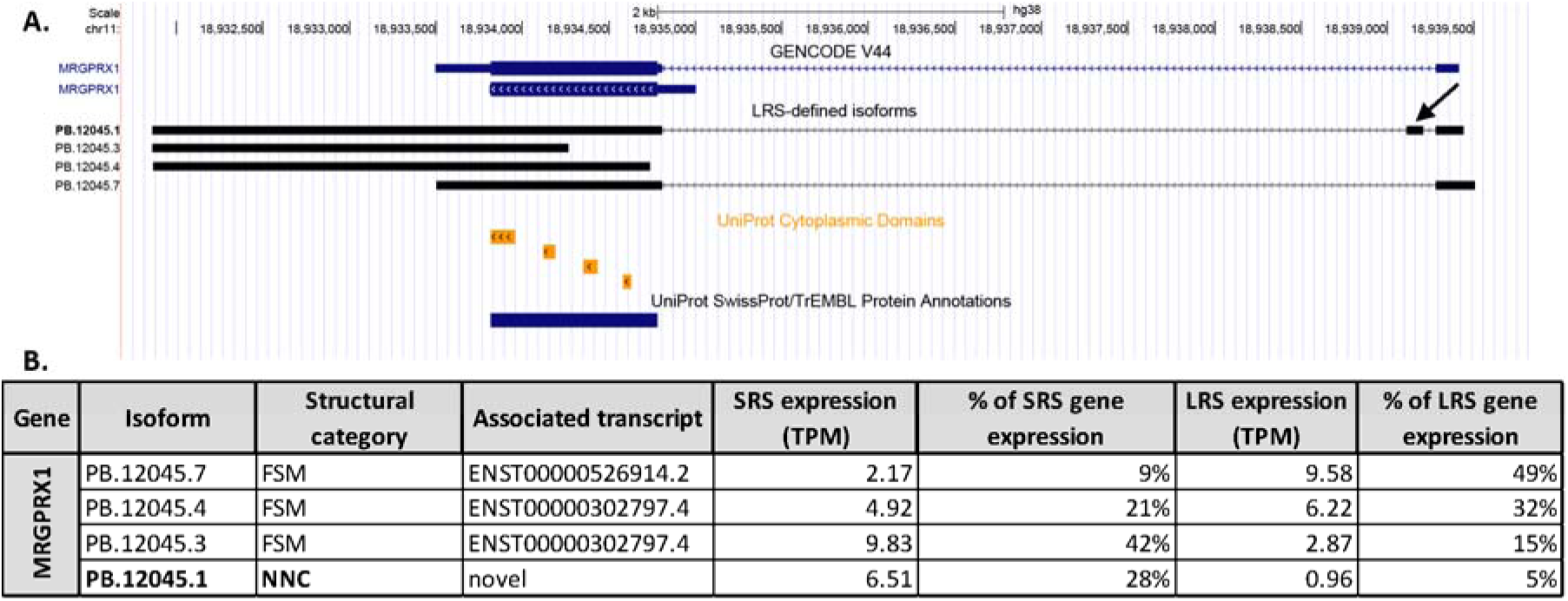
Isoforms of MRGPRX1 detected with long read sequencing. **A.** The isoforms of MRGPRX1 previously annotated in the GENCODE V44 library and the 4 LRS-identified MRGPRX1 isoforms. The isoform ID in bold indicates a novel isoform. An arrow shows the novel exon identified in isoform PB.12045.1. The UniProt cytoplasmic domains and protein annotation is additionally shown. **B.** A table of the expression level (measured as TPM) for each isoform detected with LRS showing the percentage of total gene expression measured with SRS and LRS for each isoform.

Three of these were FSM and 1 was a novel isoform utilizing novel splice-sites to include a 93bp exon in the 5’ end of the transcript. The canonical *MRGPRX1* transcript consists of 2 exons where exon 1 encodes the 5’UTR. The novel exon identified in the *MRGPRX1* isoform was upstream of the canonical start site, consequently, the open reading frame sequence of the novel isoform was predicted to be identical to the canonical FSM. An upstream in-frame stop codon was annotated in the inserted sequence (bp 177-179), however, using IRESpy no potential IRES were predicted in the 5’UTR sequence of the novel PB.12045.1 isoform. The novel isoform was identified in 2 out of 3 of the organ donors with LRS, and when mapping the SRS to the LRS-defined transcriptome, the isoform makes up 28% of the expression of the *MRGPRX1* gene (**Figure 8B**).

Sixteen novel isoforms of *HNRNPK* were identified in the hDRG with LRS, with no FSM or ISM isoforms (**Figure 9A**). Upon further inspection, the coding regions of the novel *HNRNPK* isoforms were consistent with the start and end-sites of the canonical isoform, ENST00000376263.8, however the novel isoforms consistently lacked an exon observed in all annotated isoforms (exon 14).

**Figure 9:**
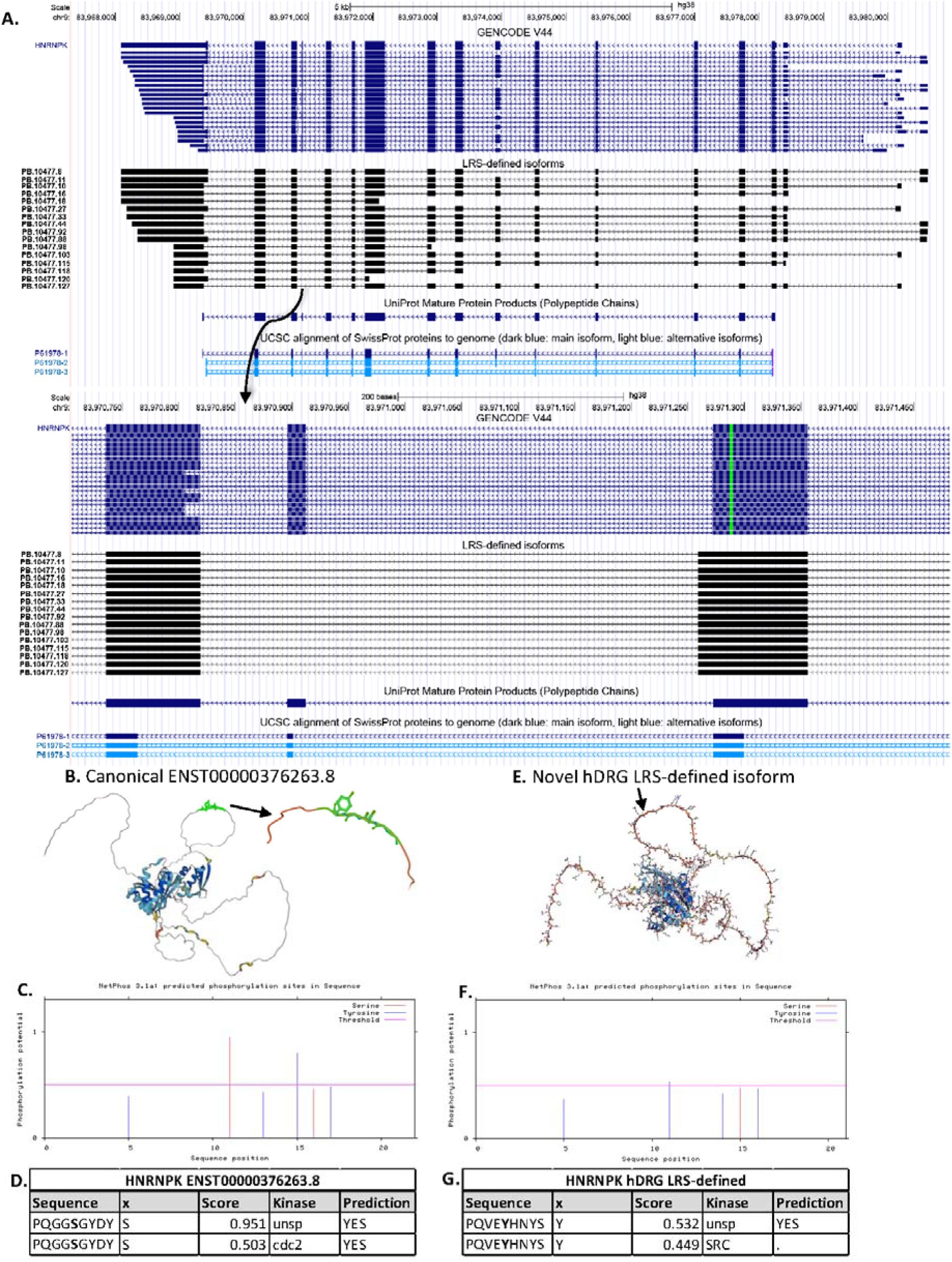
Isoforms of HNRNPK detected with long read sequencing. **A.** The isoforms of HNRNPK previously annotated in the GENCODE V44 library and the 16 novel LRS-identified isoforms are shown along with the UniProt canonical protein annotation, as well as the UCSC protein isoforms. A detailed view of the deletion of exon 14 and extension of exon 13 is shown. **B.** The protein structure for the canonical isoform, ENST00000376263.8, of HNRNPK is illustrated with AlphaFold, highlighting the 6aa region which is excised in the previously unannotated isoform. **C.** Predicted phosphorylation sites for serine and tyrosine in the canonical isoform are annotated with NetPhos3.1. A cutoff of 0.5 for phosphorylation potential is imposed. **D.** A table of the predicted serine-phosphorylation site in the hnRNP K canonical isoform in the region of the 6aa sequence shows a cdc2-phosphorylation site. **E.** The protein structure for the novel hDRG HNRNPK isoform predicted with AlphaFold v2.3.2, indicating the 5aa inserted region. **F.** Predicted phosphorylation sites for serine and tyrosine in the inserted novel region of the hDRG isoform annotated with NetPhos3.1 with a cutoff of 0.5 for phosphorylation potential. **G.** The predicted tyrosine-phosphorylation site in the inserted 5aa region most likely utilized by SRC.

Furthermore, exon 13 was found to contain an additional sequence of 13bp in the 3’end of the exon. The deletion of exon 14 and elongation of exon 13 caused a deletion of 6aa (GGSGYD), replaced by 5aa (VEYHN), while the remaining coding sequence was conserved. The structure of the primary canonical hnRNP K isoform was shown in AlphaFold, and the deletion site identified in an outlying loop structure (**Figure 9B**). Using NetPhos3.1, the deletion was predicted to remove a serine-phosphorylation site favored by cdc2 (**Figure 9C, 9D**). The novel isoform of hnRNP K with the deletion/insertion at exons 13 and 14 was modeled in AlphaFold, showing minimal structural variation apart from the change to the loop structure (**Figure 9E**).

The insertion was predicted to introduce a tyrosine-phosphorylation site with potential phosphorylation site for SRC (**Figure 9F, 9G**). The insertion/deletion was characterized as novel and unannotated in the GENCODE V44 database as well as when using the NCBI Nucleotide Blast. However, upon searching the inserted amino acid sequence, including the 8aa prior and after the sequence insertion, using the NCBI Protein Blast tool, a 100% match is identified in human proteome as hnRNP K isoform CRA_b (GenBank: EAW62676.1) (Venter et al., 2001). The hnRNPK CRA_b isoform matches the hDRG-identified hnRNP K aa sequence, apart from the last 5aa. The 5’ end of the hDRG-identified hnRNP K sequence matches the 6aa sequence of the canonical human hnRNP K sequence.

## Discussion

Our work provides a thorough overview of the isoform variants in the hDRG with a focus on splicing. Our analysis provides a resource that can be used to examine splicing across most genes that are expressed in this tissue. One of our key findings is the relatively high number of novel mRNA isoforms that are expressed in the hDRG. This could be a product of more diverse splicing in neurons or other cell types in the hDRG, or it could be a result of a paucity of data that would allow for annotation of unique splicing events that occur in this tissue. Given that previous studies suggest complex splicing patterns in the human nervous system, we favor the latter hypothesis as an explanation for our findings. Nevertheless, our findings provide new insight into gene expression patterns in the hDRG that may be very important for understanding pain and peripheral neuropathies.

We obtained high quality LRS data, which facilitated the characterization of isoforms in hDRG. Detection of short form *IFNAR2* isoform reflected results presented in other human tissues (Pasquesi et al., 2023). Furthermore, we independently validated the detection of an isoform of *WNK1* identified by Sapio et al., containing an exon not previously annotated in the GENCODE V44 library. We showed a possible functional implication of the previously unannotated exon in *WNK1*, as it was predicted to insert a sequence within the coding region with potential to interact with GSK3 kinase. Previous research has indicated the nociceptor-specific HSN2 exon in *WNK1* interacts with GSK3 to promote neurite outgrowth in primary cortical neurons of mice (Shimizu & Shibuya, 2022). Sapio et al. found that the novel exon was most commonly expressed in isoforms with the HSN2 exon, consequently we can hypothesize that the novel exon may play a role in the WNK1-HSN2/GSK3 pathway of neurite outgrowth (Sapio et al., 2023).

Novel isoforms of *SPP1*, *ASIC3*, *MRGPRX1* and *HNRNPK* were identified, utilizing splice-sites to include or exclude regions of transcripts. In *ASIC3* and *MRGPRX1* the utilization of novel splice-junctions facilitated changes to the 5’UTR of the transcripts. The significance of the 5’UTR in translation regulation has been shown in rodent and human research of the peripheral nervous system (Hudder & Werner, 2000; Ryczek et al., 2023; Song et al., 2017).

Studies in a mouse-model of Charcot-Marie-Tooth disease, showed a mutation in the 5’-UTR of nerve-specific *connexin-32* mRNA prevented function of an IRES, consequently preventing the translation of the mRNA (Hudder & Werner, 2000). Variations to the 5’-UTR, including IRES sequences, of human *OPRM1* have been shown to affect binding of RBPs including hNRNP K and PCBP1, consequently affecting protein synthesis in human neuroblastoma cells (Song et al., 2017). While *OPRM1* is detected at the RNA level, and functionally expressed in hDRG, we did not detect the transcript in LRS, likely because it is lowly expressed as shown in many bulk RNA sequencing experiments from hDRG tissue (Moy et al., 2020; North et al., 2019; Ray et al., 2018; Ray et al., 2023). We hypothesize there are changes to the protein structure and expression of osteopontin, ASIC3, and MRGPRX1 following the utilization of novel splice-junctions as observed using LRS.

In hDRG, novel isoforms of *HNRNPK* illustrated the immense functional implication of LRS results. All *HNRNPK* isoforms identified in hDRG were predicted to lack an annotated serine-phosphorylation site in favor of introducing a novel tyrosine-phosphorylation site most probably targeted by SRC. The inclusion of this new phosphorylation site introduces a mechanism for alternative regulation of translation of transcripts bound by the hnRNP K protein. Translation regulation in DRG neurons plays a critical role in the development of chronic pain in preclinical models (Khoutorsky & Price, 2018; Yousuf et al., 2021). Therefore, new mechanisms of translation regulation emerging either from novel splice isoforms of RNA binding proteins like hnRNP K or novel mRNA variants with change to the 5’ UTR could be important new contributors to chronic pain mechanisms. Loss of function mutations in the HNRNPK gene lead to Au-Kline syndrome, a rare disorder with distinct facial features and neurological deficits (Au et al., 1993). Interesting, the syndrome also features very high pain thresholds and dysautonomia (Au et al., 2018), suggesting a fundamental function of this gene in the peripheral nervous system that should be explored in more detail in future studies.

### Limitations and future directions

Our work is primarily limited by the sample size used for this research. The LRS-defined transcriptome was created with data from 3 organ donor samples and validated using SRS from 3 independent donor samples, however, the results are vulnerable to population variability. The rate of interindividual variation in isoform expression patterns are unknown, however, transcriptome-wide differences related to age, sex and pathology history have previously been shown and consequently should not be underestimated (North et al., 2019; Palmer et al., 2021; Ray et al., 2022). Future studies will address these concerns through increased sample size, facilitating the consideration of confounding factors in the analysis. Furthermore, while *in silico* validation was included in the analysis performed in this study, future work will address validation with a secondary technique, such as RNAscope or qPCR. The depth of LRS is a limitation illustrated by the absence of detection of transcripts expressed in hDRG with other sequencing technologies including *SCN9A* and *OPRM1*. The Sequel II LRS technology had limited sequencing coverage capability, which has been significantly improved with new developments to the technology now offered through Revio. Future studies can be designed to address splicing patterns in lowly expressed genes by utilizing new technologies as well as limiting the number of samples sequenced on 1 chip. Furthermore, the role of splicing in the development of chronic pain is hypothesized based on isoform discoveries for genes related to nociception, however, additional research is required.

Important questions will be answered by doing LRS on samples recovered from organ donors with a history of neuropathic pain (Iadarola et al., 2022; Renthal et al., 2021).

## Conflict of interest

The authors declare no conflicts of interest.

## Funding

This work was supported by NIH grants U19NS130608 and R01NS065926 to TJP.

## Acknowledgements

The authors are grateful to the organ donors and their families for their gift.

## Notes

### Competing Interest Statement

The authors have declared no competing interest.

## References cited

Au, P. Y. B., Goedhart, C., Ferguson, M., Breckpot, J., Devriendt, K., Wierenga, K., Fanning, E., Grange, D. K., Graham, G. E., Galarreta, C., Jones, M. C., Kini, U., Stewart, H., Parboosingh, J. S., Kline, A. D., Innes, A. M., & Care for Rare Canada, C. (2018). Phenotypic spectrum of Au-Kline syndrome: a report of six new cases and review of the literature. Eur J Hum Genet, 26(9), 1272–1281. 10.1038/s41431-018-0187-2

Au, P. Y. B., Innes, A. M., & Kline, A. D. (1993). Au-Kline Syndrome. In M. P. Adam, J. Feldman, G. M. Mirzaa, R. A. Pagon, S. E. Wallace, L. J. H. Bean, K. W. Gripp, & A. Amemiya (Eds.), GeneReviews(®). University of Washington, Seattle Copyright © 1993-2023, University of Washington, Seattle. GeneReviews is a registered trademark of the University of Washington, Seattle. All rights reserved.

Barragan-Iglesias, P., Franco-Enzastiga, U., Jeevakumar, V., Shiers, S., Wangzhou, A., Granados-Soto, V., Campbell, Z. T., Dussor, G., & Price, T. J. (2020). Type I Interferons Act Directly on Nociceptors to Produce Pain Sensitization: Implications for Viral Infection-Induced Pain. J Neurosci, 40(18), 3517–3532. 10.1523/JNEUROSCI.3055-19.2020

Bhuiyan, S. A., Xu, M., Yang, L., Semizoglou, E., Bhatia, P., Pantaleo, K. I., Tochitsky, I., Jain, A., Erdogan, B., Blair, S., Cat, V., Mwirigi, J. M., Sankaranarayanan, I., Tavares-Ferreira, D., Green, U., McIlvried, L. A., Copits, B. A., Bertels, Z., Del Rosario, J. S., Renthal, W. (2023). Harmonized cross-species cell atlases of trigeminal and dorsal root ganglia. bioRxiv. 10.1101/2023.07.04.547740

Blom, N., Gammeltoft, S., & Brunak, S. (1999). Sequence and structure-based prediction of eukaryotic protein phosphorylation sites1 1Edited by F. E. Cohen. Journal of Molecular Biology, 294(5), 1351–1362. 10.1006/jmbi.1999.3310

Blom, N., Sicheritz-Pontén, T., Gupta, R., Gammeltoft, S., & Brunak, S. (2004). Prediction of post-translational glycosylation and phosphorylation of proteins from the amino acid sequence. Proteomics, 4(6), 1633–1649. 10.1002/pmic.200300771

Donaldson, L. F., & Beazley-Long, N. (2016). Alternative RNA splicing: contribution to pain and potential therapeutic strategy. Drug Discovery Today, 21(11), 1787–1798. 10.1016/j.drudis.2016.06.017

Graveley, B. R. (2001). Alternative splicing: increasing diversity in the proteomic world. Trends in Genetics, 17(2), 100–107. 10.1016/S0168-9525(00)02176-4

Hu, T., Chitnis, N., Monos, D., & Dinh, A. (2021). Next-generation sequencing technologies: An overview. Human Immunology, 82(11), 801–811. 10.1016/j.humimm.2021.02.012

Hudder, A., & Werner, R. (2000). Analysis of a Charcot-Marie-Tooth Disease Mutation Reveals an Essential Internal Ribosome Entry Site Element in the Connexin-32 Gene*. Journal of Biological Chemistry, 275(44), 34586–34591. 10.1074/jbc.M005199200

Hulse, R. P., Beazley-Long, N., Hua, J., Kennedy, H., Prager, J., Bevan, H., Qiu, Y., Fernandes, E. S., Gammons, M. V., Ballmer-Hofer, K., Gittenberger de Groot, A. C., Churchill, A. J., Harper, S. J., Brain, S. D., Bates, D. O., & Donaldson, L. F. (2014). Regulation of alternative VEGF-A mRNA splicing is a therapeutic target for analgesia. Neurobiol Dis, 71, 245–259. 10.1016/j.nbd.2014.08.012

Iadarola, M. J., Sapio, M. R., & Mannes, A. J. (2022). Be in it for the Long Haul: A Commentary on Human Tissue Recovery Initiatives. J Pain, 23(10), 1646–1650. 10.1016/j.jpain.2022.04.009

Jumper, J., Evans, R., Pritzel, A., Green, T., Figurnov, M., Ronneberger, O., Tunyasuvunakool, K., Bates, R., Žídek, A., Potapenko, A., Bridgland, A., Meyer, C., Kohl, S. A. A., Ballard, A. J., Cowie, A., Romera-Paredes, B., Nikolov, S., Jain, R., Adler, J., Hassabis, D. (2021). Highly accurate protein structure prediction with AlphaFold. Nature, 596(7873), 583–589. 10.1038/s41586-021-03819-2

Khoutorsky, A., & Price, T. J. (2018). Translational Control Mechanisms in Persistent Pain. Trends Neurosci, 41(2), 100–114. 10.1016/j.tins.2017.11.006

Lee, P. T., Chao, P. K., Ou, L. C., Chuang, J. Y., Lin, Y. C., Chen, S. C., Chang, H. F., Law, P. Y., Loh, H. H., Chao, Y. S., Su, T. P., & Yeh, S. H. (2014). Morphine drives internal ribosome entry site-mediated hnRNP K translation in neurons through opioid receptor-dependent signaling. Nucleic Acids Res, 42(21), 13012–13025. 10.1093/nar/gku1016

Li, Y., North, R. Y., Rhines, L. D., Tatsui, C. E., Rao, G., Edwards, D. D., Cassidy, R. M., Harrison, D. S., Johansson, C. A., Zhang, H., & Dougherty, P. M. (2018). DRG Voltage-Gated Sodium Channel 1.7 Is Upregulated in Paclitaxel-Induced Neuropathy in Rats and in Humans with Neuropathic Pain. J Neurosci, 38(5), 1124–1136. 10.1523/JNEUROSCI.0899-17.2017

Lipscombe, D., & Lopez Soto, E. J. (2019). Alternative splicing of neuronal genes: new mechanisms and new therapies. Current Opinion in Neurobiology, 57, 26–31. 10.1016/j.conb.2018.12.013

Lischka, A., Lassuthova, P., Çakar, A., Record, C. J., Van Lent, J., Baets, J., Dohrn, M. F., Senderek, J., Lampert, A., Bennett, D. L., Wood, J. N., Timmerman, V., Hornemann, T., Auer-Grumbach, M., Parman, Y., Hübner, C. A., Elbracht, M., Eggermann, K., Geoffrey Woods, C., Kurth, I. (2022). Genetic pain loss disorders. Nature Reviews Disease Primers, 8(1). 10.1038/s41572-022-00365-7

Liu, S., Kang, W.-J., Abrimian, A., Xu, J., Cartegni, L., Majumdar, S., Hesketh, P., Bekker, A., & Pan, Y.-X. (2021). Alternative Pre-mRNA Splicing of the Mu Opioid Receptor Gene, OPRM1: Insight into Complex Mu Opioid Actions. Biomolecules, 11(10), 1525. https://www.mdpi.com/2218-273X/11/10/1525

Liu, Y., Cao, C., Huang, X.-P., Gumpper, R. H., Rachman, M. M., Shih, S.-L., Krumm, B. E., Zhang, S., Shoichet, B. K., Fay, J. F., & Roth, B. L. (2023). Ligand recognition and allosteric modulation of the human MRGPRX1 receptor. Nature Chemical Biology, 19(4), 416–422. 10.1038/s41589-022-01173-6

Lopes, I., Altab, G., Raina, P., & de Magalhães, J. P. (2021). Gene Size Matters: An Analysis of Gene Length in the Human Genome. Front Genet, 12, 559998. 10.3389/fgene.2021.559998

Moy, J. K., Hartung, J. E., Duque, M. G., Friedman, R., Nagarajan, V., Loeza-Alcocer, E., Koerber, H. R., Christoph, T., Schroder, W., & Gold, M. S. (2020). Distribution of functional opioid receptors in human dorsal root ganglion neurons. Pain. 10.1097/j.pain.0000000000001846

Nguyen, M. Q., von Buchholtz, L. J., Reker, A. N., Ryba, N. J. P., & Davidson, S. (2021). Single-nucleus transcriptomic analysis of human dorsal root ganglion neurons. eLife, 10, e71752. 10.7554/eLife.71752

North, R. Y., Li, Y., Ray, P., Rhines, L. D., Tatsui, C. E., Rao, G., Johansson, C. A., Zhang, H., Kim, Y. H., Zhang, B., Dussor, G., Kim, T. H., Price, T. J., & Dougherty, P. M. (2019). Electrophysiological and transcriptomic correlates of neuropathic pain in human dorsal root ganglion neurons. Brain, 142(5), 1215–1226. 10.1093/brain/awz063

Palmer, C. R., Liu, C. S., Romanow, W. J., Lee, M. H., & Chun, J. (2021). Altered cell and RNA isoform diversity in aging Down syndrome brains. Proc Natl Acad Sci U S A, 118(47). 10.1073/pnas.2114326118

Pan, Q., Shai, O., Lee, L. J., Frey, B. J., & Blencowe, B. J. (2008). Deep surveying of alternative splicing complexity in the human transcriptome by high-throughput sequencing. Nature Genetics, 40(12), 1413–1415. 10.1038/ng.259

Pardo-Palacios, F. J., Arzalluz-Luque, A., Kondratova, L., Salguero, P., Mestre-Tomás, J., Amorín, R., Estevan-Morió, E., Liu, T., Nanni, A., McIntyre, L., Tseng, E., & Conesa, A. (2023). SQANTI3: curation of long-read transcriptomes for accurate identification of known and novel isoforms. bioRxiv, 2023.2005.2017.541248. 10.1101/2023.05.17.541248

Pasquesi, G. I. M., Allen, H., Ivancevic, A., Barbachano-Guerrero, A., Joyner, O., Guo, K., Simpson, D. M., Gapin, K., Horton, I., Nguyen, L., Yang, Q., Warren, C. J., Florea, L. D., Bitler, B. G., Santiago, M. L., Sawyer, S. L., & Chuong, E. B. (2023). Regulation of interferon signaling by transposon exonization. Cold Spring Harbor Laboratory. 10.1101/2023.09.11.557241

Raj, B., & Blencowe, Benjamin J. (2015). Alternative Splicing in the Mammalian Nervous System: Recent Insights into Mechanisms and Functional Roles. Neuron, 87(1), 14–27. 10.1016/j.neuron.2015.05.004

Ray, P., Torck, A., Quigley, L., Wangzhou, A., Neiman, M., Rao, C., Lam, T., Kim, J. Y., Kim, T. H., Zhang, M. Q., Dussor, G., & Price, T. J. (2018). Comparative transcriptome profiling of the human and mouse dorsal root ganglia: an RNA-seq-based resource for pain and sensory neuroscience research. Pain, 159(7), 1325–1345. 10.1097/j.pain.0000000000001217

Ray, P. R., Shiers, S., Caruso, J. P., Tavares-Ferreira, D., Sankaranarayanan, I., Uhelski, M. L., Li, Y., North, R. Y., Tatsui, C., Dussor, G., Burton, M. D., Dougherty, P. M., & Price, T. J. (2022). RNA profiling of human dorsal root ganglia reveals sex differences in mechanisms promoting neuropathic pain. Brain, 146(2), 749–766. 10.1093/brain/awac266

Ray, P. R., Shiers, S., Caruso, J. P., Tavares-Ferreira, D., Sankaranarayanan, I., Uhelski, M. L., Li, Y., North, R. Y., Tatsui, C., Dussor, G., Burton, M. D., Dougherty, P. M., & Price, T. J. (2023). RNA profiling of human dorsal root ganglia reveals sex differences in mechanisms promoting neuropathic pain. Brain, 146(2), 749–766. 10.1093/brain/awac266

Renthal, W., Chamessian, A., Curatolo, M., Davidson, S., Burton, M., Dib-Hajj, S., Dougherty, P. M., Ebert, A. D., Gereau, R. W. t., Ghetti, A., Gold, M. S., Hoben, G., Menichella, D. M., Mercier, P., Ray, W. Z., Salvemini, D., Seal, R. P., Waxman, S., Woolf, C. J., Price, T. J. (2021). Human cells and networks of pain: Transforming pain target identification and therapeutic development. Neuron, 109(9), 1426–1429. 10.1016/j.neuron.2021.04.005

Ryczek, N., Łyś, A., & Makałowska, I. (2023). The Functional Meaning of 5’UTR in Protein-Coding Genes. Int J Mol Sci, 24(3). 10.3390/ijms24032976

Sapio, M. R., King, D. M., Staedtler, E. S., Maric, D., Jahanipour, J., Kurochkina, N. A., Manalo, A. P., Ghetti, A., Mannes, A. J., & Iadarola, M. J. (2023). Expression pattern analysis and characterization of the hereditary sensory and autonomic neuropathy 21A (HSAN2A) gene with no lysine kinase (WNK1) in human dorsal root ganglion. Experimental Neurology, 114552. 10.1016/j.expneurol.2023.114552

Sharon, D., Tilgner, H., Grubert, F., & Snyder, M. (2013). A single-molecule long-read survey of the human transcriptome. Nat Biotechnol, 31(11), 1009–1014. 10.1038/nbt.2705

Shekarabi, M., Girard, N., Rivière, J. B., Dion, P., Houle, M., Toulouse, A., Lafrenière, R. G., Vercauteren, F., Hince, P., Laganiere, J., Rochefort, D., Faivre, L., Samuels, M., & Rouleau, G. A. (2008). Mutations in the nervous system--specific HSN2 exon of WNK1 cause hereditary sensory neuropathy type II. J Clin Invest, 118(7), 2496–2505. 10.1172/jci34088

Shimizu, M., & Shibuya, H. (2022). WNK1/HSN2 mediates neurite outgrowth and differentiation via a OSR1/GSK3β-LHX8 pathway. Sci Rep, 12(1), 15858. 10.1038/s41598-022-20271-y

Song, K. Y., Choi, H. S., Law, P.-Y., Wei, L.-N., & Loh, H. H. (2017). Post-Transcriptional Regulation of the Human Mu-Opioid Receptor (MOR) by Morphine-Induced RNA Binding Proteins hnRNP K and PCBP1. Journal of Cellular Physiology, 232(3), 576–584. 10.1002/jcp.25455

Song, Y., Wang, Z.-Y., Luo, J., Han, W.-C., Wang, X.-Y., Yin, C., Zhao, W.-N., Hu, S.-W., Zhang, Q., Li, Y.-Q., & Cao, J.-L. (2023). CWC22-Mediated Alternative Splicing of Spp1 Regulates Nociception in Inflammatory Pain. Neuroscience. 10.1016/j.neuroscience.2023.10.006

Stamm, S., Ben-Ari, S., Rafalska, I., Tang, Y., Zhang, Z., Toiber, D., Thanaraj, T. A., & Soreq, H. (2005). Function of alternative splicing. Gene, 344, 1–20. 10.1016/j.gene.2004.10.022

Stark, R., Grzelak, M., & Hadfield, J. (2019). RNA sequencing: the teenage years. Nature Reviews Genetics, 20(11), 631–656. 10.1038/s41576-019-0150-2

Su, Y., Fan, L., Shi, C., Wang, T., Zheng, H., Luo, H., Zhang, S., Hu, Z., Fan, Y., Dong, Y., Yang, J., Mao, C., & Xu, Y. (2021). Deciphering Neurodegenerative Diseases Using Long-Read Sequencing. Neurology, 97(9), 423–433. 10.1212/wnl.0000000000012466

Tardaguila, M., De La Fuente, L., Marti, C., Pereira, C., Pardo-Palacios, F. J., Del Risco, H., Ferrell, M., Mellado, M., Macchietto, M., Verheggen, K., Edelmann, M., Ezkurdia, I., Vazquez, J., Tress, M., Mortazavi, A., Martens, L., Rodriguez-Navarro, S., Moreno-Manzano, V., & Conesa, A. (2018). SQANTI: extensive characterization of long-read transcript sequences for quality control in full-length transcriptome identification and quantification. Genome Research, 28(3), 396–411. 10.1101/gr.222976.117

Tavares-Ferreira, D., Shiers, S., Ray, P. R., Wangzhou, A., Jeevakumar, V., Sankaranarayanan, I., Cervantes, A. M., Reese, J. C., Chamessian, A., Copits, B. A., Dougherty, P. M., Gereau, R. W., Burton, M. D., Dussor, G., & Price, T. J. (2022). Spatial transcriptomics of dorsal root ganglia identifies molecular signatures of human nociceptors. Science Translational Medicine, 14(632), eabj8186. doi:10.1126/scitranslmed.abj8186

Venter, J. C., Adams, M. D., Myers, E. W., Li, P. W., Mural, R. J., Sutton, G. G., Smith, H. O., Yandell, M., Evans, C. A., Holt, R. A., Gocayne, J. D., Amanatides, P., Ballew, R. M., Huson, D. H., Wortman, J. R., Zhang, Q., Kodira, C. D., Zheng, X. H., Chen, L., Zhu, X. (2001). The Sequence of the Human Genome. Science, 291(5507), 1304–1351. doi:10.1126/science.1058040

Wang, E. T., Sandberg, R., Luo, S., Khrebtukova, I., Zhang, L., Mayr, C., Kingsmore, S. F., Schroth, G. P., & Burge, C. B. (2008). Alternative isoform regulation in human tissue transcriptomes. Nature, 456(7221), 470–476. 10.1038/nature07509

Wang, J., & Gribskov, M. (2019). IRESpy: an XGBoost model for prediction of internal ribosome entry sites. BMC Bioinformatics, 20(1), 409. 10.1186/s12859-019-2999-7

Wang, Y., Liu, J., Huang, B., Xu, Y.-M., Li, J., Huang, L.-F., Lin, J., Zhang, J., Min, Q.-H., Yang, W.-M., & Wang, X.-Z. (2015). Mechanism of alternative splicing and its regulation. Biomedical Reports, 3(2), 152–158. 10.3892/br.2014.407

Wangzhou, A., McIlvried, L. A., Paige, C., Barragan-Iglesias, P., Shiers, S., Ahmad, A., Guzman, C. A., Dussor, G., Ray, P. R., Gereau, R. W. t., & Price, T. J. (2020). Pharmacological target-focused transcriptomic analysis of native vs cultured human and mouse dorsal root ganglia. Pain, 161(7), 1497–1517. 10.1097/j.pain.0000000000001866

Weyn-Vanhentenryck, S. M., Feng, H., Ustianenko, D., Duffié, R., Yan, Q., Jacko, M., Martinez, J. C., Goodwin, M., Zhang, X., Hengst, U., Lomvardas, S., Swanson, M. S., & Zhang, C. (2018). Precise temporal regulation of alternative splicing during neural development [Article]. Nature Communications, 9(1), Article 2189. 10.1038/s41467-018-04559-0

Xu, C., Prete, M., Webb, S., Jardine, L., Stewart, B., Hoo, R., He, P., & Teichmann, S. A. (2023). Automatic cell type harmonization and integration across Human Cell Atlas datasets. Cold Spring Harbor Laboratory. 10.1101/2023.05.01.538994

Xu, J., Lu, Z., Xu, M., Pan, L., Deng, Y., Xie, X., Liu, H., Ding, S., Hurd, Y. L., Pasternak, G. W., Klein, R. J., Cartegni, L., Zhou, W., & Pan, Y.-X. (2014). A Heroin Addiction Severity-Associated Intronic Single Nucleotide Polymorphism Modulates Alternative Pre-mRNA Splicing of the μ Opioid Receptor Gene <EM>OPRM1</EM> via hnRNPH Interactions. The Journal of Neuroscience, 34(33), 11048–11066. 10.1523/jneurosci.3986-13.2014

Yang, T. H., Wang, C. Y., Tsai, H. C., & Liu, C. T. (2021). Human IRES Atlas: an integrative platform for studying IRES-driven translational regulation in humans. Database (Oxford), 2021. 10.1093/database/baab025

Yousuf, M. S., Shiers, S. I., Sahn, J. J., & Price, T. J. (2021). Pharmacological Manipulation of Translation as a Therapeutic Target for Chronic Pain. Pharmacol Rev, 73(1), 59–88. 10.1124/pharmrev.120.000030

Yu, H., Usoskin, D., Nagi, S. S., Hu, Y., Kupari, J., Bouchatta, O., Cranfill, S. L., Gautam, M., Su, Y., Lu, Y., Wymer, J., Glanz, M., Albrecht, P., Song, H., Ming, G.-L., Prouty, S., Seykora, J., Wu, H., Ma, M., Luo, W. (2023). Single-Soma Deep RNA Sequencing of Human Dorsal Root Ganglion Neurons Reveals Novel Molecular and Cellular Mechanisms Underlying Somatosensation. bioRxiv, 2023.2003.2017.533207. 10.1101/2023.03.17.533207

Zeng, P., Tian, Z., Han, Y., Zhang, W., Zhou, T., Peng, Y., Hu, H., & Cai, J. (2022). Comparison of ONT and CCS sequencing technologies on the polyploid genome of a medicinal plant showed that high error rate of ONT reads are not suitable for self-correction. Chinese Medicine, 17(1). 10.1186/s13020-022-00644-1

Zhang, P., Perez, O. C., Southey, B. R., Sweedler, J. V., Pradhan, A. A., & Rodriguez-Zas, S. L. (2021). Alternative Splicing Mechanisms Underlying Opioid-Induced Hyperalgesia. Genes, 12(10), 1570. 10.3390/genes12101570

